# Alternate subunit assembly diversifies the function of a bacterial toxin

**DOI:** 10.1101/624130

**Authors:** Casey Fowler, Gabrielle Stack, Xuyao Jiao, Maria Lara-Tejero, Jorge E. Galán

## Abstract

Bacterial toxins with an AB_5_ architecture are central to bacterial pathogenesis. Functionally diverse and evolutionarily distant AB_5_ toxins adopt synonymous structures in which a discrete domain of the toxin’s active (A) subunit is inserted into a ring-like platform comprised of five delivery (B) subunits. *Salmonella* Typhi, the cause of typhoid fever, produces an unusual A_2_B_5_ toxin known as typhoid toxin, a major virulence factor. Here, we report that upon infection of human cells, *S*. Typhi produces two forms of typhoid toxin that have distinct delivery components but share common active subunits. We demonstrate that the two typhoid toxins exhibit substantially different trafficking properties, elicit markedly different effects when administered to laboratory animals, and are expressed in response to different regulatory mechanisms and distinct metabolic cues. Collectively, these results indicate that the evolution of two typhoid toxin variants has conferred functional versatility to this virulence factor. More broadly, this study reveals a new paradigm in toxin biology and suggests that the evolutionary expansion of AB_5_ toxins was likely fueled by the remarkable plasticity inherent to their structural design coupled to the functional versatility afforded by the combination of homologous toxin components.

## Main Text

*Salmonella enterica* serovar Typhi (*S*. Typhi) is the etiological agent of typhoid fever, a major global health problem^1,2^. An assortment of evidence indicates that typhoid toxin is responsible for some of the more severe symptoms of typhoid fever^3–5^. Compared to other AB-type toxins, typhoid toxin is highly unusual in that two A subunits, CdtB, a DNAse, and PltA, an ADP-ribosyltransferase, associate with a single pentameric B subunit, PltB, resulting in a unique A_2_B_5_ architecture^4^. This unusual composition appears to be the result of typhoid toxin’s remarkable evolutionary history, during which two classes of AB-type toxins – cytolethal distending toxins (CDTs) and pertussis-like toxins - amalgamated to produce a single toxin^6^. A structural and biochemical investigation into the evolution of typhoid toxin revealed that this class of toxins exhibits remarkable plasticity in that heterologous co-expression of various combinations of homologs from other bacteria produced active typhoid-toxin-like complexes^6^.

Given the observed plasticity in toxin assembly, we were intrigued by the observation that *S*. Typhi encodes a genomic islet containing a locus with significant sequence similarity to a pertussis-like toxin (Fig. 1a, Extended Data Fig. 1). However, although the gene for the delivery B subunit (*sty1364*) appears to encode a full-length protein, the gene encoding the active A subunit (*sty1362*) contains several stop codons and a ~200 bp C-terminal deletion, indicating that it is a pseudogene no longer encoding a functional protein. Therefore, *sty1364*, a *pltB* homolog, appears to encode an orphan B subunit, which we hypothesized may associate with PltA and CdtB to produce an alternative form of typhoid toxin (Fig. 1a). In testing this hypothesis, we found that *sty1364* exhibited a similar pattern of expression to the genes encoding other components of typhoid toxin^3,7,8^ in that, although undetectable under standard laboratory conditions, it was induced > 100-fold following *S*. Typhi infection of cultured human epithelial cells (Fig. 1b). Similarly, *sty1364* expression was strongly induced in *S*. Typhi grown in TTIM, a growth medium that mimics some aspects of the intracellular environment and is permissive for typhoid toxin expression *in vitro*^8^ (Fig. 1c). Using affinity chromatography coupled to LC-MS/MS and western blot analysis we readily detected an interaction between Sty1364 and both CdtB and PltA after *S*. Typhi growth in TTIM medium as well as after infection of cultured epithelial cells (Figs. 1d, 1e, Supplementary Table 1). Importantly, Sty1364 did not interact with CdtB in the absence of PltA, indicating that similar to the holotoxin assembled with PltB, Sty1364 forms a complex with CdtB only through its interaction with PltA (Figs 1d, 1e). Sty1364 therefore appears to represent a bona fide typhoid toxin component and thus we have renamed it PltC. Although most AB_5_ toxins employ a homopentameric B subunit ^9^, pertussis toxin is comprised of heteropentameric B subunit assembled from different but structurally related components^10,11^. However, we did not detect PltB in samples affinity purified from a tag present in PltC, indicating that typhoid toxin does not exhibit a single heteromeric B subunit architecture but rather it is assembled in two alternative configurations with PltB or PltC as its homopentameric subunit (Fig. 1d-f). We also observed an increase in the amount of PltB that co-immunoprecipitated with CdtB in a Δ*pltC* strain compared to wild type, indicating that, in the absence of PltC, more PltB-containing toxin is assembled thus suggesting that the two delivery subunits compete for their association to the active subunits (Figs. 1e, 1f). Collectively, these data indicate that upon infection of human cells *S*. Typhi assembles two distinct typhoid toxins with the same enzymatic “A” subunits but distinct delivery platforms or “B” subunits (Fig. 1a).

**Fig. 1.**
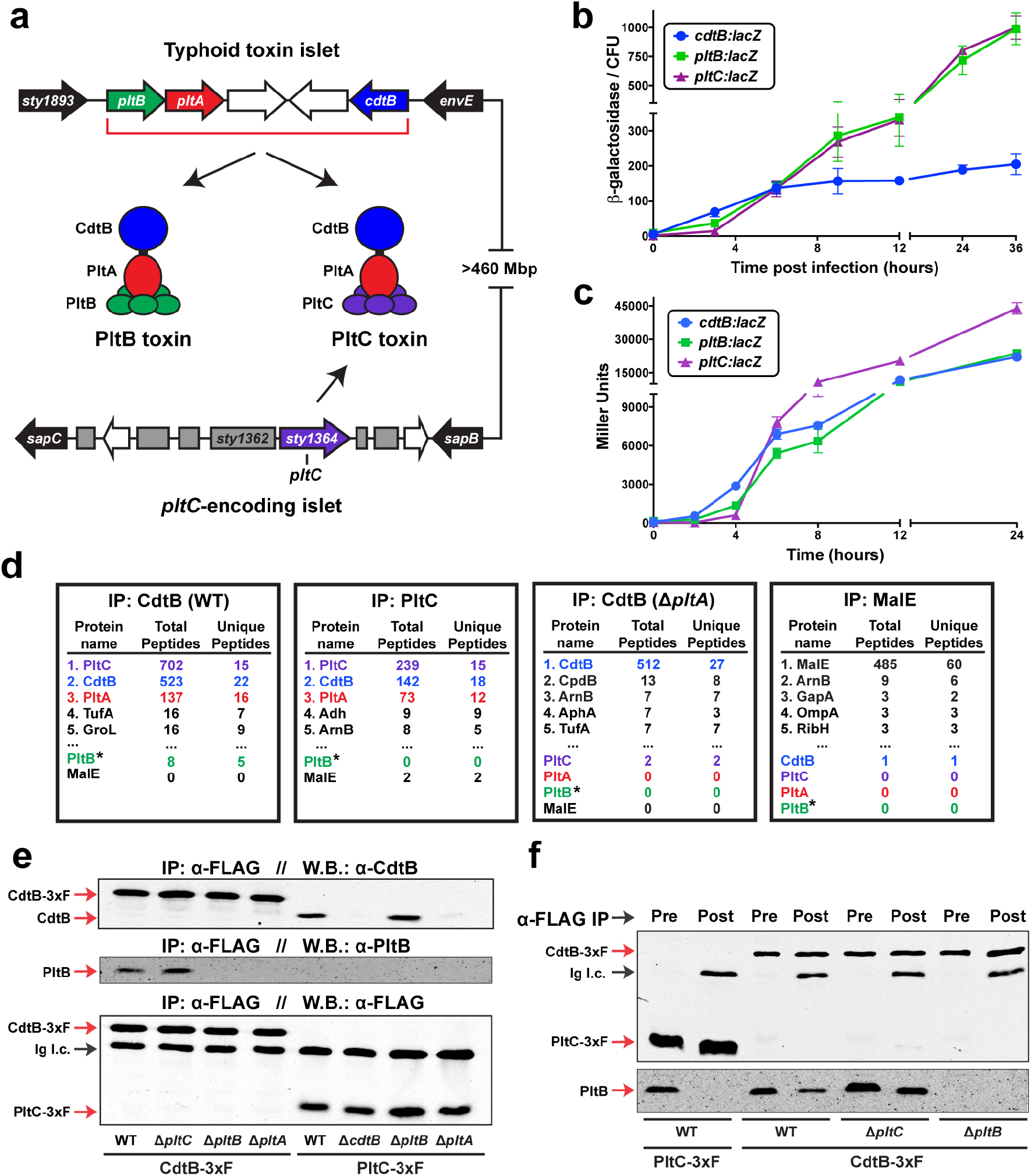
*S*. Typhi produces two distinct typhoid toxins that employ common active subunits but different delivery subunits. (**a**) Illustration of the *S*. Typhi typhoid toxin genomic locus, as well as a distant locus that encodes *pltC* (*sty1364*), an orphan pertussis-like toxin delivery subunit that exhibits homology to *pltB*. (**b** and **c**) Expression levels of *pltC*, *pltB* and *cdtB* over time upon encountering conditions that stimulate typhoid toxin gene expression. The β-galactosidase activity, normalized to the number of bacteria, from *pltC:lacZ, pltB:lacZ* and *cdtB:lacZ S*. Typhi reporter strains was measured at the indicated time points following infection of Henle-407 cells (**b**) or growth in TTIM medium (**c**). Values indicate the mean +/- S.D. for three independent samples. (**d**) Interaction of PltC with PltA/CdtB in *S*. Typhi grown in TTIM. Four *S*. Typhi strains encoding *cdtB*-3xFLAG, *pltC*-3xFLAG, *cdtB*-3xFLAG/Δ*pltA*, or *malE*-3xFLAG (as a control) were grown in TTIM to induce typhoid toxin expression. Cell lysates were immunoprecipitated with an anti-FLAG antibody and interacting proteins were identified using LC/MS/MS. For each sample the numbers of peptides and unique peptides are shown for the five most abundant *S*. Typhi proteins recovered and for all typhoid toxin subunits, which are color-coded to coincide with the schematic shown in panel (**a**). * Due to its small size and the nature of its amino acid sequence the identification of PltB using our LC/MS/MS protocol is inefficient even in purified typhoid toxin preparations. (**e**) *S*. Typhi produces both PltB- and PltC-typhoid toxins within infected human cells. Henle-407 cells were infected with *S*. Typhi wild type or the indicated mutant strains encoding 3xFLAG epitope-tagged CdtB or PltC (as indicated) and 24 hs post-infection the interaction of the indicated toxin components were probed by co-immunoprecipitation and western blot analysis. (**f**) PltB forms a complex with CdtB, but not with PltC. The indicated strains featuring 3xFLAG epitope tagged CdtB or PltC were grown in TTIM for 24 hs to induce typhoid toxin expression and the interaction of the indicated toxin components in cell lysates were probed by co-immunoprecipitation and western blot analysis. Ig. l. c.: Immunoglobulin light chain detected by the secondary antibody.

To assess the function of the typhoid toxin assembled with PltC we examined the ability of the purified toxin to intoxicate cultured cells as measured by cell cycle arrest at G2/M due to the DNA damage inflicted by the CdtB subunit (Figs. 2a, 2b)^3,7,12^. We found that PltC-typhoid toxin was able to intoxicate cultured epithelial cells in a similar fashion to the PltB version of the toxin although with a higher (~7 fold) EC50. Interestingly, however, cultured cells infected with a strain that lacks *pltC* (exclusively producing PltB-typhoid toxin) were intoxicated in a manner indistinguishable to cells infected with wild type *S*. Typhi, although cells infected with a Δ*pltB* mutant strain (exclusively producing PltC-typhoid toxin) did not exhibit detectable signs of intoxication (Fig. 2c). The observations that PltC-typhoid toxin is produced to significant levels during infection (Fig. 1e) and is able to intoxicate when directly applied to cultured cells, but it does not intoxicate during bacterial infection suggested that the two alternative forms of the toxin might differ in their delivery mechanisms after their synthesis by intracellular *S*. Typhi. We have previously shown that following its production, typhoid toxin is secreted from the bacterial periplasm into the lumen of the Salmonella-containing vacuole (SCV) by a specialized protein secretion mechanism involving a specialized muramidase that enables the toxin to cross to the trans side of the PG layer, from where it can be released by various membrane-active agonists such as bile salts or anti-microbial peptides^13^. We found that both forms of typhoid toxin are released from the bacteria using this same mechanism (Extended Data Fig. 2). It has been shown that after its secretion into the lumen of the SCV, typhoid toxin is packaged into vesicle transport intermediates that carry the toxin to the extracellular space, a process that is orchestrated by interactions of its B subunit PltB with specific luminal receptors^3,14^. Therefore we examined whether differences in receptor specificity between PltB and PltC may lead to differences in their intracellular transport pathways after bacterial infection. We infected cultured cells with wild-type *S*. Typhi or isogenic mutants carrying deletions in *pltC, pltB*, or both, and monitored the formation of CdtB-containing transport carriers using an immunofluorescence assay (Figs. 2d, 2e). We found that toxin carriers were absent in cells infected with the Δ*pltB* strain although they were readily detected in cells infected with the Δ*pltC* mutant. These results indicate that the formation of the transport carriers is strictly dependent on PltB, presumably because the PltC version of typhoid toxin does not engage the sorting receptor. In fact, the level of transport carriers was measurably increased in cells infected with the Δ*pltC S*. Typhi mutant, an indication that the absence of PltC leads to the assembly of higher levels of export-competent PltB-containing typhoid toxin. Collectively, these data indicate that the two typhoid toxins differ significantly with respect to their ability to engage the sorting receptors within the lumen of the SCV leading to substantial differences in the export of the toxins to the extracellular space, a pre-requisite for intoxication after bacterial infection of cultured cells. These results also suggest that, during infection, the two toxins may exert their function in different environments and may target different cells.

**Fig. 2.**
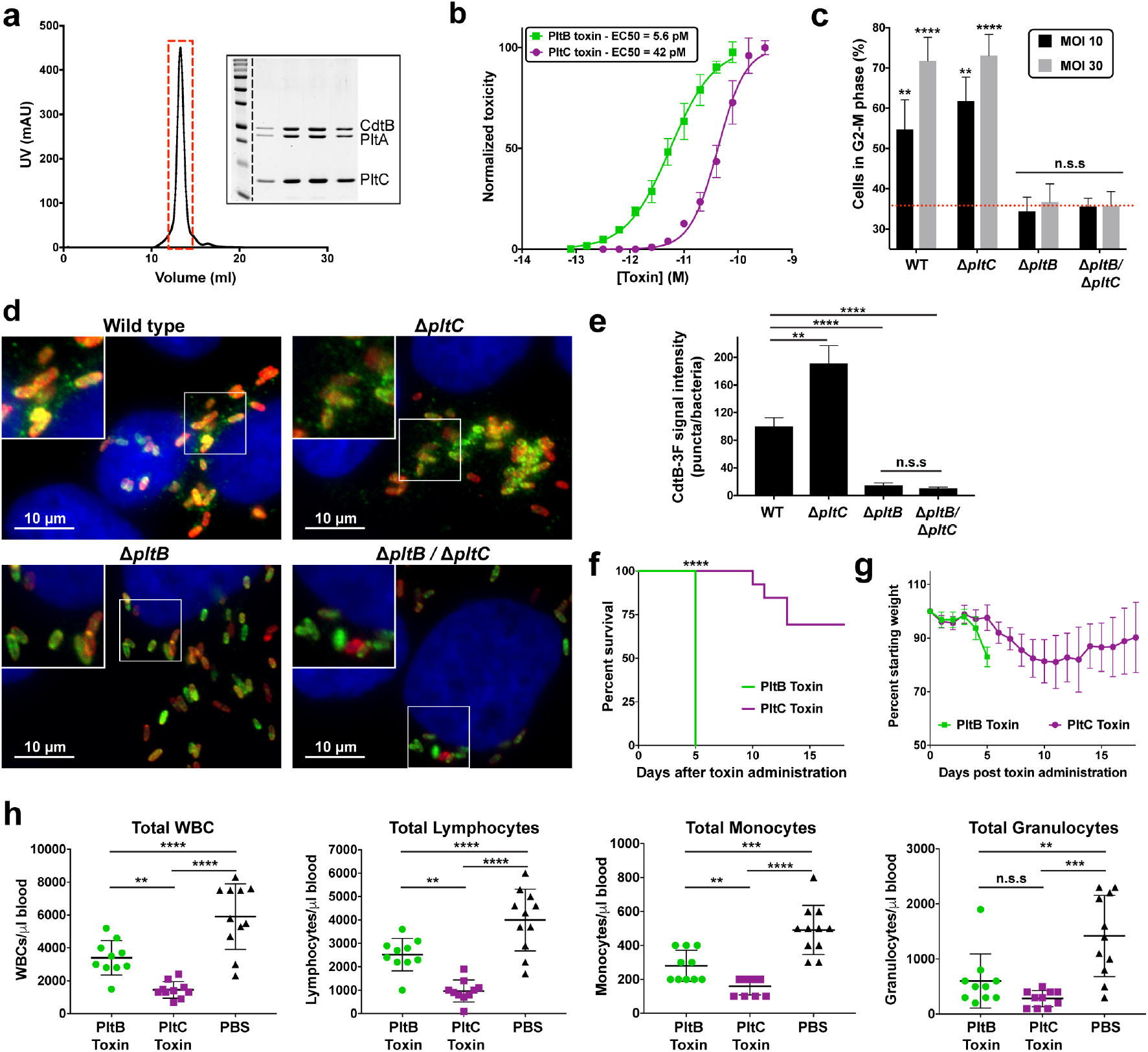
The PltB- and PltC-typhoid toxins exhibit significantly different trafficking properties and elicit different effects *in vivo*. (**a**) Purification of the PltC toxin. Gel filtration chromatography and SDS-PAGE/Coomassie (inset) blue analyses of purified PltC-typhoid toxin. (**b**) PltC-typhoid toxin elicits G2/M cell cycle arrest in human epithelial cells. Purified PltB- or PltC-typhoid toxins were added to the culture medium of Henle-407 cells at the indicated concentrations and 48 hs after, cells were fixed and analyzed by flow cytometry to evaluate toxicity as indicated in Materials and Methods. The data shown are the mean normalized toxicity +/- S.D. for three independent experiments. (**c**) PltC-typhoid toxin does not induce G2/M arrest in *S*. Typhi-infected cells. Henle-407 cells were infected with the indicated strains at a multiplicity of infection (MOI) of 10 or 30, as indicated, and 48 hs post-infection cells were collected and the percentage of cells in G2/M phase was determined as described for panel (**b**). Mean values +/- S.D. are shown for three independent experiments assayed in duplicate (6 total samples). Asterisks denote statistically significant levels G2/M cell cycle arrest compared to mock infected cells (red dotted line) as determined by unpaired two-tailed t-tests. (**d**) and (**e**) PltC-typhoid toxin is not packaged into vesicle transport carriers. Henle-407 cells were infected with the indicated *S*. Typhi strains encoding 3x-FLAG epitope-tagged CdtB and 48 hs postinfection the cells were fixed and stained with DAPI (blue), α-FLAG (green), and α-*S*. Typhi LPS (red) antibodies. Typhoid toxin-containing export vesicles, which appear as green puncta (**d**), were quantified by image analysis (**e**) as indicated in Materials and Methods. Values are from at least 30 images (~100 infected cells) taken in two independent experiments and represent the mean relative ratios +/- S.E.M. Asterisks denote the statistical significance of the indicated pairwise comparisons determined using unpaired two-tailed t-tests. (**f-h**) The PltB- and PltC-typhoid toxins elicit different effects when administered to mice. Highly purified preparations of PltB- (2 μg) or PltC-typhoid toxins (10 μg) were administered to C57BL/6 mice. For one group of mice, their survival (**f**) and body weight (mean +/- S.D.) (**g**) was recorded at the indicated times. The remaining mice were sacrificed at four days post-toxin administration and a blood sample was collected and analyzed to quantify the indicated cell types (mean +/- S.D.) (**h**). WBCs, white blood cells. The Mantel-Cox test was used for statistical analysis of mouse survival and unpaired two-tailed t-tests were used to statistically compare the indicated samples for the blood analysis. For all panels, **** p<0.0001, *** p<0.001, ** p<0.01, * p<0.05, n.s.s. not statistically significant.

To gain insight into potential differences between the activities of the two forms of typhoid toxin, we evaluated the consequences of systemically administering to C57BL/6 mice highly purified preparations of PltB- or PltC-typhoid toxins. We found that, although administering 2 μg of PltB-typhoid toxin was sufficient to kill all mice tested within five days, all mice administered 10 μg of PltC-Typhoid toxin (~5-fold more), survived for at least 10 days and the majority survived the full course of the experiment (Fig. 2f). Furthermore, in contrast to PltB-typhoid toxin^15^, mice receiving the PltC-typhoid toxin showed no neurological symptoms, but did lose weight and showed signs of malaise and lethargy, although these symptoms were delayed and less severe than those observed in the PltB-typhoid toxin treated animals (Fig. 2g). Peak toxicity for PltC-typhoid toxin treated mice was observed ~8-13 days post-administration, after which the majority of treated animals fully recovered. Interestingly, despite eliciting fewer and milder overt symptoms compared the PltB-typhoid toxin treated mice, PltC-typhoid toxin caused a significantly greater reduction in the numbers of total white blood cells, lymphocytes and monocytes (Figure 2h). Collectively, these data indicate that the two typhoid toxins preferentially target different cells/tissues. Therefore producing two toxins variants confers functional versatility to typhoid toxin, presumably enabling *S*. Typhi to expand the spectrum of host cell targets that it can engage.

Given the substantially different functional properties exhibited by the two forms of typhoid toxin, we reasoned that *S*. Typhi might have evolved regulatory mechanisms to preferentially produce the different forms of the toxin under different conditions. Expression of the typhoid toxin locus genes (i. e. *pltB, pltA and cdtB*) is controlled by the PhoP/PhoQ (PhoPQ) two-component regulatory system^8^. However, we found that in the absence of PhoPQ, *pltC* was robustly expressed during bacterial infection indicating that, despite exhibiting a similar intracellular expression pattern to the other typhoid toxin genes, the regulation of *pltC* expression must be distinct (Extended Data Figs. 3, 4). To decipher its regulatory network, we applied FAST-INseq^8^ to screen for *S*. Typhi genes that influence *pltC* expression within infected cultured cells (Fig. 3a). Cultured epithelial cells were infected with a library of *S*. Typhi transposon mutants that encode a GFP reporter of *pltC* expression and, using fluorescence activated cell sorting (FACS), bacterial mutants that expressed *pltC* were separated from those that did not. Transposon insertion site sequencing (INseq)^16–18^ was then used to identify transposon-disrupted genes that were over-represented in the bacterial population that did or did not express *pltC*, thus identifying candidate genes required for the regulation of *pltC* expression (Fig. 3b and Supplementary Table 2). Notably, the most significantly enriched mutants that did not express *pltC* were insertions within *ssrA* and ssrB, which encode a two-component regulatory system (Extended Data Table 1)^19–21^. This system is the master regulator of the expression of a type III protein secretion system encoded within its pathogenicity island 2, an essential *Salmonella* virulence factor that, like typhoid toxin, is selectively expressed by intracellular bacteria^22–24^. *S*. Typhi strains carrying deletion mutations in *ssrA/ssrB*, showed drastically reduced levels of *pltC* expression following infection of cultured epithelial cells, confirming the observations in the genetic screen (Fig. 3C, Extended Data Fig. 4). In contrast, absence of SsrA/SsrB had a negligible effect on *pltB* expression. Collectively, these results suggest that the SsrA/SsrB two-component system is the principal activator of *pltC* expression during infection and indicate that the production of the two typhoid toxin delivery subunits is controlled by different, intracellularly-induced, global regulatory networks. Our screen also identified several mutants that led to increased *pltC* expression. Among these mutants were insertions within all of the genes required for the biosynthesis of the cofactor biotin (Fig. 3b and Extended Data Table 2), which based on previous studies^8^, are likely to affect expression of *pltC* indirectly, by preventing the expansion of a population of cytosolic bacteria (i. e. located outside of the *Salmonella* containing vacuole) unable to express typhoid toxin. These results therefore suggest that, like the other typhoid toxin genes, *pltC* expression also requires the specific environment of the *Salmonella* containing vacuole. We also found that insertions within genes required for purine biosynthesis resulted in increased *pltC* expression, while insertions within pyrimidine biosynthesis genes had the opposite effect (Fig. 3b, Extended Data Tables 1, 2). This is particularly noteworthy since purine biosynthesis mutants result in reduced expression of pltB^8^. Follow up experiments exploring the expression of *pltB* and *pltC* in cultured cells infected with isogenic purine (Δ*purM*) or pyrimidine (Δ*pyrC*) biosynthesis mutants confirmed the results of our screen and demonstrated that, in intracellular *S*. Typhi, purine limitation favors *pltC* expression while pyrimidine limitation favors *pltB* expression (Figs. 3d and 3e). Collectively, these results demonstrate that the relative expression levels of the two typhoid toxin delivery subunits are substantially altered in response to low nucleotide concentrations, and suggest that different nutrient availability may serve as a cue that enables *S*. Typhi to adjust the balance of the two forms of typhoid toxin it produces within a given environment (Fig. 3f).

**Fig. 3.**
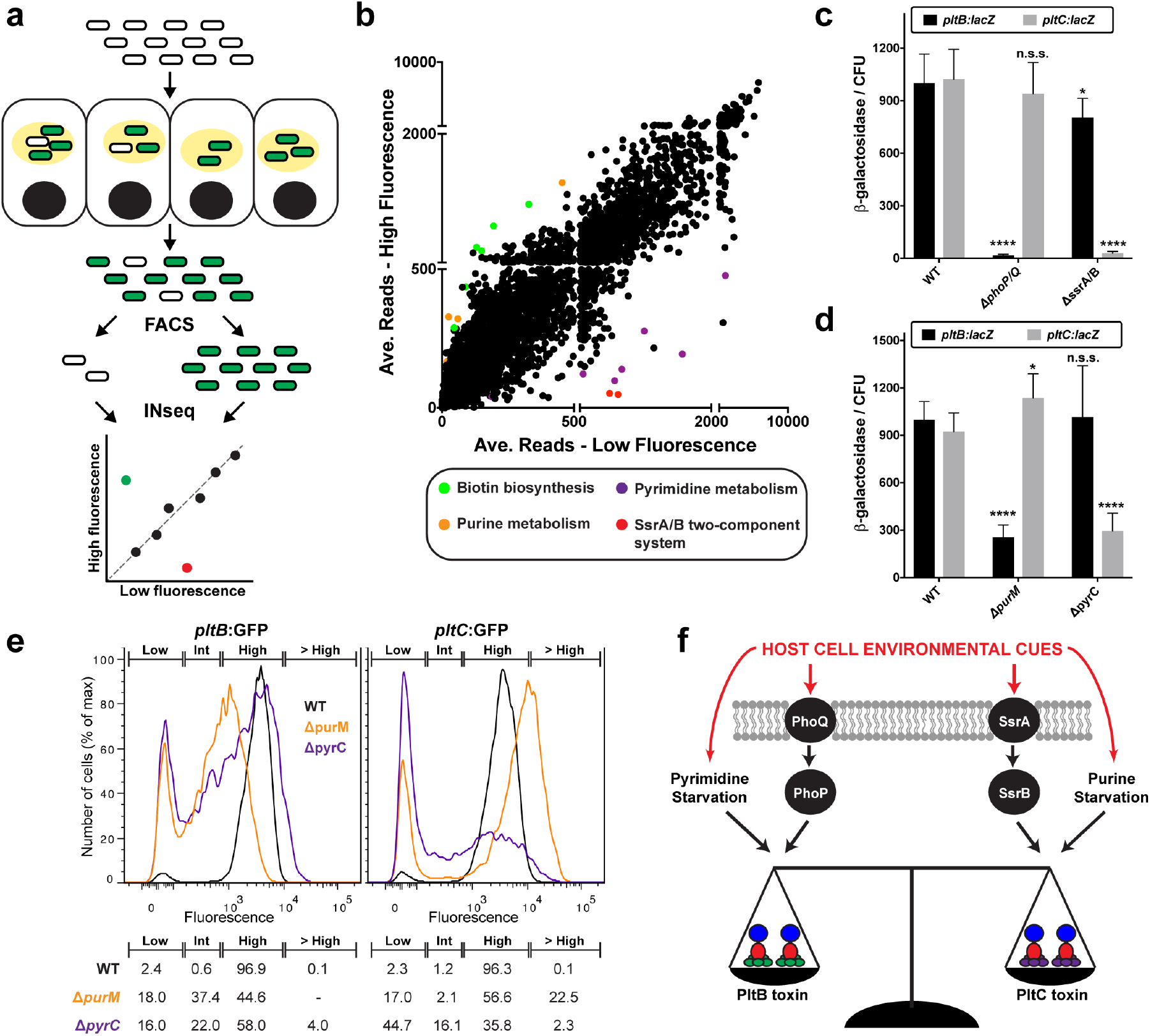
Distinct regulatory mechanisms and metabolic cues control the expression of the different typhoid toxin delivery subunits. (**a**) Schematic representation of the FAST-INseq genetic screen used to identify *S*. Typhi genes that influence *pltC* expression in infected host cells. A large library of random transposon mutants was generated in the *S*. Typhi *pltC:gfp* strain and used to infect Henle-407 cells. Sixteen hours post-infection the bacteria were isolated and sorted by FACS into pools exhibiting high and low levels of GFP fluorescence. The populations of transposon insertions in these two pools were then determined using INseq in order to identify mutants that stimulate (green dot in plot) or reduce (red dot) *pltC* expression during infection. (**b**) Overview of the results of the FAST-INseq screen. Plot shows the normalized numbers of sequencing reads of transposon insertions within each *S*. Typhi gene in the high fluorescence pool versus the low fluorescence pool. (**c** and **d**) Expression levels of *pltB:lacZ* and *pltC:lacZ* reporters in infected human cells for wild-type *S*. Typhi (WT) and the indicated deletion mutant strains. Henle-407 cells were infected with the indicated strains for 24 hours, after which the β-galactosidase activity from bacterial lysates was measured and normalized to the numbers of CFU recovered. Values indicate mean values +/- S.D. for six independent determinations taken over two separate experiments. Asterisks denote statistically significant differences relative to the corresponding wild-type sample determined using unpaired two-tailed t-tests. **** p<0.0001, * p<0.05, n.s.s. not statistically significant. (**e**) Flow cytometry analysis of *pltB:gfp* and *pltC:gfp* expression of the indicated *S*. Typhi strains 24 hours post-infection. Histograms show the GFP fluorescence intensities of individual bacteria for the indicated strains. Gates were established to show the percentage of bacteria exhibiting high, low and intermediate (int) fluorescence. The percentage of bacteria with fluorescence intensities within these gates is shown (bottom). (**f**) Overview of the identified factors that differentially affect the expression of *pltB* and *pltC* and thus are likely to be important for controlling relative abundance of the two typhoid toxins produced by *S*. Typhi upon encountering different environments during infection.

We have shown here that *S*. Typhi produces two different versions of typhoid toxin that share their enzymatic subunits but utilize alternative delivery subunits resulting is substantially different biological activities. Therefore, by assembling toxins with different targeting mechanisms, *Salmonella* Typhi may be able to target a broader array of cell types in different tissue environments. In addition to the *S*. Typhi serovar, the typhoid toxin genomic islet is sporadically present in the genomes of other *Salmonellae* that with few exceptions also encode a second B subunit ortholog (Extended Data Fig. 5) that in at least one instance genetic evidence indicate can form a functional toxin^25–27^. The distributions and genomic locations of these elements indicate that the acquisition of the core genomic islet and the second B subunit occurred in separate evolutionary events in several distinct *Salmonella* lineages, implying that producing functionally diverse typhoid toxins is a prevalent phenomenon that imparts a strong evolutionary advantage to ecologically divergent *Salmonellae* (Extended Data Fig. 5). More broadly, we have also identified other pathogens that encode orphan B subunits that are homologous to components of an AB_5_ toxin located elsewhere in their genomes, suggesting that the assembly of alternate toxins may be a more general feature of AB_5_ toxins (Extended Data Fig. 6). For example, in addition to the locus encoding pertussis toxin, *Bordetella pertussis* encodes two orphan orthologs of its B subunits at a distant genome location. Under certain conditions, these proteins have been reported to be co-synthesized and co-secreted with other pertussis toxin components^28^. The results shown here also suggest that the evolutionary expansion of the AB_5_ class of toxins was likely fueled by the plasticity inherent to their structural design coupled with the functional versatility that can be achieved through combining homologous toxin components.

## Acknowledgments

We thank members of the Galán laboratory for careful review of this manuscript. C.C.F. was supported in part by a Postdoctoral Fellowship from the Canadian Institutes of Health Research (CIHR). This work was supported by National Institute of Allergy and Infectious Diseases grant AI079022 (to J.E.G.).

## Methods

### Bacterial Strains and Cell lines

*S*. Typhi strains employed in this study were derived from the wild-type isolate ISP2825^29^ and were constructed using standard recombinant DNA and allelic exchange procedures using the *E. coli* β-2163 Δ*nic35* as the conjugative donor strain ^30^. Strains were routinely cultured in LB broth at 37°C. For *in vitro* growth assays that employed typhoid toxin inducing growth conditions (TTIM) a previously described chemically defined growth medium was used ^8^ that was based on N minimal medium ^31^. All experiments using cultured cells were conducted using the Henle-407 human epithelial cell line, which was obtained from the Roy Curtiss library collection. Cells were cultured in Dulbecco’s modified Eagle medium (DMEM, GIBCO) supplemented with 10% Fetal Bovine Serum (FBS) at 37°C with 5% CO_2_ in a humidified incubator. This cell line was routinely tested for mycoplasma contamination using a Mycoplasma Detection Kit (SouthernBiotech, Cat# 13100-01).

### *Salmonella* Typhi infections

To infect Henle-407 cells, overnight cultures of *S*. Typhi were diluted 1/20 into fresh LB containing 0.3 M NaCl and grown to an OD_600_ of 0.9. Cells were infected for 1 hour in Hank’s balanced salt solution (HBSS) at the indicated multiplicity of infection (MOI). Cells were then washed three times with HBSS and incubated in culture medium containing 100 μg/ml gentamycin to kill extracellular bacteria. After 1 hour, cells were washed and fresh medium was added containing 5 μg/ml gentamycin to avoid repeated cycles of reinfection.

### β-galactosidase assays

For *in vitro* grown samples, overnight cultures were washed two times with TTIM, diluted 1/20 into fresh TTIM and grown for the indicated amount of time at 37°C at which point 10 or 20 μl of the culture was added to 90 μl of permeabilization buffer (100 mM Na_2_HPO_4_, 20 mM KCl, 2 mM MgSO_4_, 0.8 mg/ml hexadecyltrimethylammonium bromide [CTAB], 0.4 mg/ml sodium deoxycholate, 5.4 μL/ml β-mercaptoethanol) and assayed as described below. For samples collected from infected cells, 3×10^5^ Henle-407 cells were plated in 6-well plates and grown for 24 hours prior to infection with the indicated strains. Following infections, cells were washed two times with PBS, released from the plates using dilute trypsin and pelleted by centrifugation for 5 minutes at 150 x g. Cells were then lysed in 0.1% sodium deoxycholate (in PBS) supplemented with 100 μg/ml DNase I and the lysate was centrifuged at 5,000 x g for 5 minutes to pellet the bacteria. The bacteria were re-suspended in PBS, a small aliquot of which was diluted to calculate the total number CFU recovered. The remainder of the bacteria were re-suspended in 100 μl of permeabilization buffer and assayed as described below. Assays were conducted at 24 hours post infection (hpi) unless otherwise indicated. β-galactosidase assays were conducted using a modified version of the protocol developed by Miller ^32 8^. Briefly, samples were permeabilized for 20 minutes at room temperature in the buffer described above. 600 μl of substrate buffer (60 mM Na_2_HPO_4_, 40 mM NaH_2_PO_4_, 2.7 μL/mL β-mercaptoethanol, 1 mg/ml ONPG [o-nitrophenyl-β-D-galactopyranoside, Sigma]) was then added to initiate the reactions. Once the samples developed an obvious yellow colour, the reactions were quenched using 700 μl of 1M Na_2_CO_3_ and the reaction time was noted. Cell debris was removed by centrifugation at 20,000 x g for 5 minutes and the OD_420_ of the samples was measured. Miller units were calculated as: (1000 * OD_420_) / (reaction time [minutes] * culture volume assayed [ml] * OD_600_ [culture]).

### Co-immunoprecipitation experiments to identify interactions between toxin components

To identify interaction partners for the various typhoid toxin subunits, *S*. Typhi bacterial cell lysates were immunoprecipitated using C-terminal 3xFLAG epitope-tagged CdtB or PltC (tags were incorporated at the native genomic locus) as indicated and the eluates were analyzed by LC-MS/MS or western blot. A strain encoding a C-terminal 3xFLAG epitope-tagged version of MalE, a periplasmic protein that is expressed in TTIM, which was cloned in place of *pltC* at the *pltC* locus in the *S*. Typhi chromosome and included as a negative control for the LC-MS/MS analysis. For *in vitro* grown samples, the indicated strains were grown overnight in LB, washed twice using TTIM, diluted 1/20 into 12ml of fresh TTIM and grown overnight. Cultures were pelleted, re-suspended in lysis buffer (50 mM Tris pH 7.5, 170 mM NaCl, cOmplete mini protease inhibitors [Sigma], 40 ug/ml DNase I) and lysed using the One Shot cell disruption system (Constant Systems, Ltd). Clarified lysates were immunoprecipitated overnight at 4°C using ANTI-FLAG M2 affinity gel (Sigma). Immunoprecipitated samples were washed thoroughly using 50 mM Tris pH 7.5/170 mM NaCl/50 mM galactose/0.1% Triton X-100 and eluted using 0.1M glycine-HCl (LC-MS/MS) or SDS-PAGE loading buffer (western blot analysis). For LC-MS/MS analysis, the eluted samples were precipitated overnight at -20°C in 80% acetone and washed twice using 80% acetone. The samples were then reduced using DTT, alkylated using iodoacetamide and trypsin digested overnight. C18 column purified peptides were then analyzed by LC/MS/MS as previously described ^33^ and MS/MS scans were processed and searched using MASCOT (Matrix Science Ltd.). The resulting peptide and protein assignments were filtered to keep only those identifications with scores above extensive homology. For western blot analysis, eluted samples were run on 15% SDS-PAGE and transferred to nitrocellulose membranes. Membranes were blocked using 5% non-fat milk, incubated overnight at 4°C with the indicated primary antibody followed by a 2 hour incubation with a fluorescently conjugated α-mouse (FLAG) or α-rabbit (PltB) secondary antibody and analyzed using the Li-Cor Odyssey blot imager.

For samples isolated from infected cells, three 15 cm dishes containing a total of 7.5×10^7^ Henle-407 cells were infected at an MOI of 20 as described above using each of the indicated *S*. Typhi strains. 24 hours post-infection, cells were washed twice using PBS and collected using a cell scraper, pelleted and washed once using PBS. Cell pellets were then resuspended in lysis buffer (50 mM Tris pH 7.5, 170 mM NaCl, complete mini protease inhibitors [Sigma], 40 ug/ml DNase I, 10 mM N-ethylmaleimide) and lysed using a Branson Digital Sonifier (3 seconds on/8 seconds off, 35% amplitude, 3 minutes total). Clarified lysates were then immunoprecipitated and analyzed by western blot as described above.

### Typhoid toxin purification

Both typhoid toxins were purified according to a previously established protocol ^15^. Briefly, *pltB, pltA* and cdtB-6xHis (PltB-typhoid toxin) or *pltC, pltA* and cdtB-6xHis (PltC-typhoid toxin) were cloned into pET28a(+) vector (Novagen). *E. coli* strains carrying these expression vectors were grown to OD_600_ ~ 0.8, at which time 250 μM IPTG was added to induce the expression of the toxin genes and the cultures were grown overnight at 30°C. Bacterial cells were pelleted by centrifugation and lysed. Crude lysates were affinity purified using Nickel resin (Qiagen), followed by cation exchange chromatography using a Mono S column (Sigma-Aldrich) and finally gel filtration using a Superdex-200 column (Sigma-Aldrich). The final fractions were analyzed by SDS-PAGE to confirm purity.

### Culture cell intoxication assays

To assess typhoid toxin toxicity in cultured human cells, the number of cells arrested in G2/M (as a consequence of CdtB-mediated DNA damage) was determined using flow cytometry as previously described ^15^. For experiments using purified toxin, 2.5×10^4^ Henle-407 epithelial cells were plated in 12-well plates. Twenty four hours later the cells were washed and fresh medium containing the indicated concentrations of PltB- or PltC-typhoid toxin were added. Forty eight hours later, the cells were removed from the dishes using trypsin treatment, pelleted, re-suspended in 300 μl PBS, fixed by adding ice cold ethanol dropwise to a final concentration of 70% and incubated overnight at -20°C. Cells were then washed with PBS, re-suspended in 500 μl of PBS containing 50 μg/ml propidium iodide, 0.1 μg/ml RNase A and 0.05% Triton X-100 and incubated for 30 minutes at 37°C. Cells were then washed and re-suspend in PBS, filtered and analyzed by a flow cytometry. The DNA content of cells was determined using FlowJo (Treestar). In order to obtain EC_50_ values, the percentage of cells in the G2/M phase (% G2/M) was determined for each sample and converted to normalized toxicity by subtracting the % G2-M value observed of untreated cells and diving by (maximum % G2/M - untreated % G2/M). The maximum % G2/M value was considered to be 90% based on our experience that values above this number are not reliably observed even at saturating toxin concentrations. For experiments using *S*. Typhi infected cells, 2.5×10^4^ Henle-407 cells were plated in 12-well plates and were infected using the indicated *S*. Typhi strains at the indicated MOI values 24 hours later. Forty eight hours post-infection the cells were collected, fixed, stained and analyzed as described above.

### Typhoid toxin secretion assay

To assess the mechanism of secretion for the PltB- and PltC typhoid toxins, an *in vitro* toxin secretion assay was employed to determine whether toxin secretion from Δ*pltB* (producing exclusively PltC-typhoid toxin) and Δ*pltC* (producing exclusively PltB-typhoid toxin) *S*. Typhi strains was dependent upon both TtsA and outer membrane perturbation, in accordance with the recently described mechanism of typhoid toxin secretion ^13^. The indicated strains, all of which encode 3x-FLAG epitope-tagged *cdtB* from its native genomic locus, were grown overnight in LB, washed twice using TTIM, diluted 1/20 into fresh TTIM and grown for 24 hours at 37°C to induce typhoid toxin and *ttsA* expression. The bacteria were then pelleted, washed thoroughly, and incubated for 15 minutes either in 0.075% bile salts (Sigma) in PBS or in PBS alone (mock treated). The bacteria were then pelleted and the supernatants were filtered using a 0.2 μm filter and TCA precipitated. The amount of toxin in the pellet and supernatant fractions was then determined by western blot analysis using an M2 α-FLAG antibody as described above.

### Immunofluorescence microscopy assay for typhoid toxin export

To compare the export of typhoid toxin from the *Salmonella-containing* vacuole to the extracellular space for PltB-typhoid toxin and PltC-typhoid toxin, we employed a previously established immunofluorescence-based assay that quantifies the levels typhoid toxin that are within exocytic vesicles compared to the levels associated with bacteria ^14^. Henle-407 cells plated on glass coverslips were infected with the indicated cdtB-3xFLAG epitope-tagged *S*. Typhi strains at a moiety of infection of 10. Forty eight hours post-infection, the samples were fixed in 4% paraformaldehyde and blocked using 3% BSA/0.3% triton X-100 in PBS. The coverslips were then incubated with a 1:5,000 dilution of mouse monoclonal M2 anti-FLAG antibody (Sigma) and a 1:10,000 dilution of rabbit polyclonal anti-*S*. Typhi LPS antibody (Sifin) overnight at 4°C. After thoroughly washing in PBS, samples were then stained using Alexa 488-conjugated anti-mouse and Alexa 594-conjugaed anti-rabbit antibodies and DAPI (Sigma) for 2 hours at room temperature, washed extensively using PBS, mounted on coverslips, and imaged imaged using an Eclipse TE2000 inverted microscope (Nikon) with an Andor Zyla 5.5 sCMOS camera driven by Micromanager software (https://www.micro-manager.org).

The open source software ImageJ (http://rsbweb.nih.gov/ij/) was used to quantify toxin export in images captured in random fields as described previously ^14^. Briefly, the LPS signal was used to identify the bacterial cells and the CdtB-3xFLAG signal was used to identify typhoid toxin. The typhoid toxin signal within the area associated with bacterial cells was subtracted from the total typhoid toxin signal in order to obtain the signal associated with typhoid toxin carrier intermediates. In a given field, the fluorescence intensity of the typhoid toxin signal that was associated with toxin carriers was normalized to the bacterial-associated typhoid toxin signal within the same field.

### Animal intoxication experiments

All animal experiments were conducted according to protocols approved by Yale University’s Institutional Animal Care and Use Committee. C57BL/6 mice were anesthetized with 30% w/v isoflurane in propylene glycol and 100 μl of toxin solution containing the indicated concentration of purified PltB- or PltC-typhoid toxin was administered via the retro-orbital route. Changes in behavior, weight and survival of the toxin-injected mice were closely monitored for the duration of the experiment. Blood samples were collected by cardiac puncture 4 days after toxin administration in Microtainer tubes coated with EDTA, kept at room temperature and analyzed within 2 hours after blood collection using a HESKA Veterinary Hematology System.

### FAST-INSeq screen

The FAST-INSeq screen was employed to identify *S*. Typhi genes that, when disrupted by transposon mutagenesis, lead to altered expression of a *pltC:gfp* reporter within infected Henle-407 epithelial cells. This screen was conducted as previously described ^8^ using a *pltB:gfp* reporter. Transposon mutagenesis was conducted using a mariner transposon delivered by pSB4807, which was mobilized into the *pltC:gfp S*. Typhi strain by conjugation using *E. coli* β-2163 Δ*nic35* as the donor strain. Mutants were selected by plating on LB agar plates containing 30 μg/ml chloramphenicol. A total of ~150,000 mutants were collected and multiple aliquots of this library were stocked for subsequent use in the screen. For each iteration of the screen, five 15 cm dishes were seeded with 1×10^7^ Henle-407 cells each and grown for ~24 hours prior to infection using the transposon mutant library described above. Aliquots of the inoculum used for infection were collected for subsequent INSeq preparation (inoculum pool). 18 hours post infection, the cells were washed three times with PBS, detached from plates using dilute trypsin and pelleted by centrifugation at 150 x g for 5 minutes. Cells were lysed using a 5 min incubation in 0.1% sodium deoxycholate (in PBS) supplemented with 100 μg/ml DNase I to degrade and solubilize the genomic DNA released from lysed host cells. The lysate was centrifuged at 5,000 x g for 5 minutes to isolate *S*. Typhi from the soluble cellular debris. The *S*. Typhi-containing pellet was re-suspended in PBS and further purified from cellular debris using two spins at 150 x g for 5 minutes (discarding the pellet fraction) and one spin at 5,000 x g for 5 minutes (discarding the supernatant). An aliquot of the recovered *S*. Typhi was amplified by growth in LB at 37°C and subsequently prepared for INSeq sequencing (post-infection pool) and the remainder was washed and diluted in PBS to a concentration of ~2×10^6^ bacteria/ml for FACS. A total of ~2×10^7^ *S*. Typhi mutants were sorted according to their fluorescence intensity in the GFP channel (488 nm excitation, 515/20 with 505LP emission) using a BD FACS Aria II flow cytometer. The isolated low fluorescence and high fluorescence pools were amplified by growth in LB at 37°C and subsequently prepared for INSeq sequencing. The screen was conducted using the same mutant library on two independent occasions, and the low fluorescence pools from these two sorts were pooled and re-sorted (re-sort of low fluorescence populations).

For INSeq sequencing, genomic DNA was extracted from each of the mutant pools, digested with MmeI (New England Biolabs) and barcoded samples were prepared for sequencing as described previously ^34^. The purified 121 bp DNA products containing barcodes to identify the individual mutant pools were sequenced on an Illumina HiSeq2000 system at the Yale Center for Genomic Analysis. The sequencing data were analyzed using the INSeq_pipeline_v3 package ^35^, which separated sequencing reads by pool, mapped/quantified insertions and grouped insertions by gene. For each pool, the total number of sequencing reads was normalized to be 2,176,000 (an average of 500/gene). To identify genes in which insertions were enriched in a statistically significant manner in one pool compared to another, a value of 50 (10% of the average number of reads per gene) was added to the normalized number of reads in both pools. Ratios of the log-transformed read numbers for the two pools were then calculated. Genes with values that were an average of more than two standard deviations from the mean over the two primary replicates of the screen and more than one standard deviation from the mean in all three sorts were considered to be significantly enriched.

### Analytical flow cytometry

To probe *pltB* and *pltC* expression within infected cells at the single bacterium level, we employed a previously developed flow cytometry-based assay ^8^. The indicated *pltC:gfp* and *pltB:gfp* strains carrying a plasmid driving constitutive mCherry expression were used to infect Henle-407 cells as described above. At 24 hours post infection, bacteria were isolated and prepared for flow cytometry as described above for the FAST-INSeq screen. At this time point, which was chosen to capture a state of purine/pyrimidine starvation in the purM/pyrC mutants, we find very few instances of *S*. Typhi within the host cell cytosol and thus virtually all *S*. Typhi express high levels of both *pltB* and *pltC* in the wild-type strains ^8^. For each sample the fluorescence intensity in the GFP channel (488 nm excitation, 515/20 with 505LP emission) was analyzed for at least 5,000 mCherry-positive particles (532 nm excitation, 610/20 with 600LP emission) using a BD FACS Aria II flow cytometer. All samples were prepared and analyzed by flow cytometry in parallel. High and low fluorescence populations were defined based on the peaks observed in the wild-type samples for the given reporter strain. The intermediate population was defined as having a fluorescence intensity between and the low and high gates and the “greater than high fluorescence” population was defined as having a fluorescence intensity greater than the high fluorescence peak (> 99.9% of particles in the wild-type sample).

---

**Extended Data Fig. 1.**
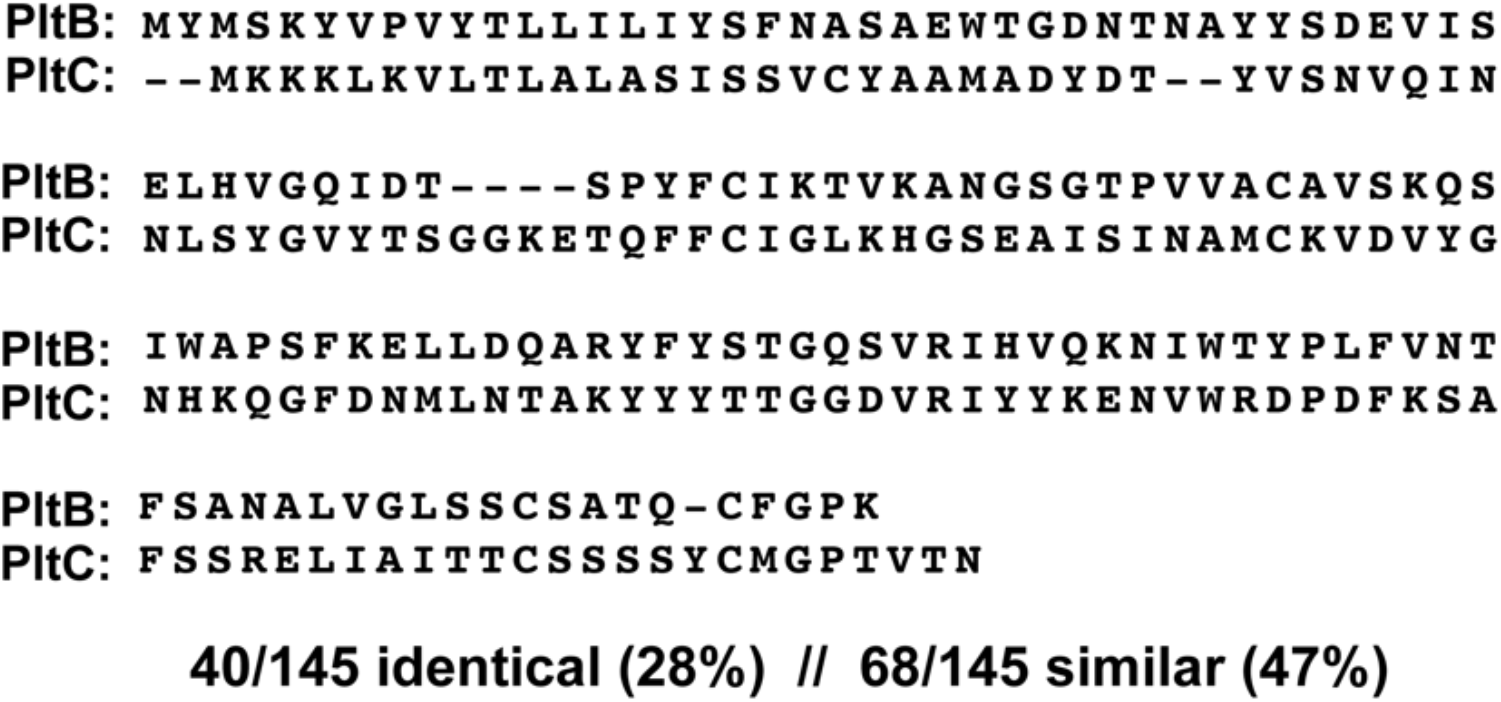
Sequence alignment of *S*. Typhi PltB and PltC. Amino acid sequence comparison between *S*. Typhi PltB and PltC (Sty1364). A pairwise global alignment was done using the EMBL-EBI Matcher program (https://www.ebi.ac.uk/Tools/psa/emboss_matcher/nucleotide.html)

**Extended Data Fig. 2.**
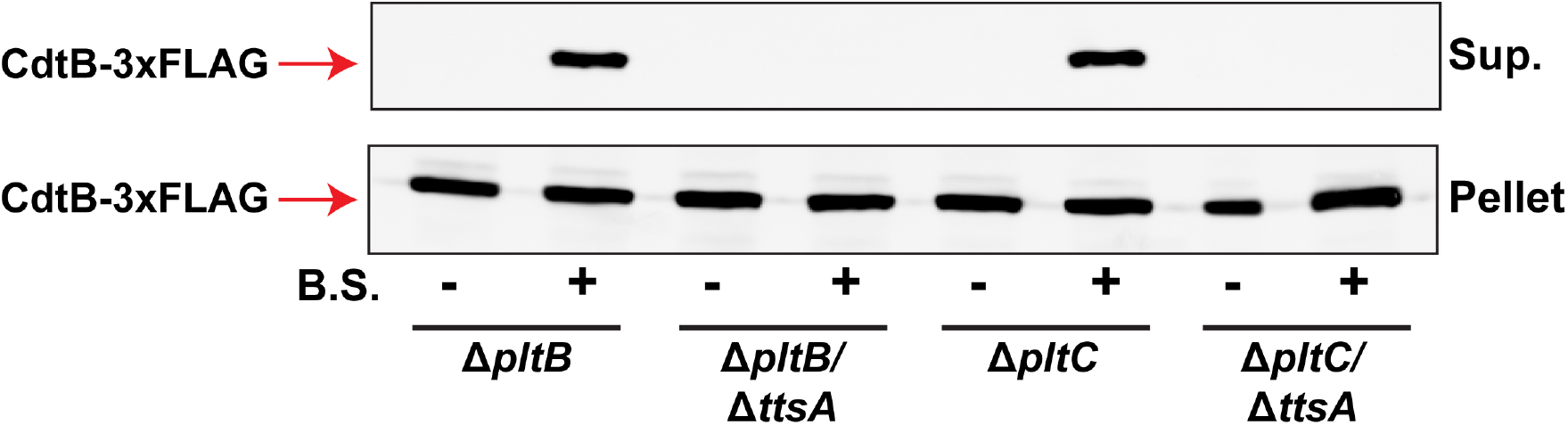
Both typhoid toxins are secreted using the same TtsA-dependent secretion mechanism. The mechanism of typhoid toxin secretion was recently described, in which the activity of the specialized muramidase TtsA enables the toxin to access the trans side of the peptidoglycan layer, after which it can be released following exposure to outer membrane perturbing agents such as bile salts ^13^. To determine whether this mechanism is utilized by both PltC-typhoid toxin and PltB-typhoid toxin, an *in vitro* secretion assay was performed ^13^. The indicated strains, all of which encode a 3xFLAG-epitope tagged CdtB, were grown for 24 hours in TTIM medium to induce typhoid toxin (and *ttsA*) expression, after which the bacteria were pelleted and washed thoroughly. The material was evenly divided between two samples, one of which was incubated in PBS containing 0.075% bile salts (B.S.) and the other in PBS only. Samples were then pelleted and the levels of CdtB-3xFLAG in the pellet fraction and in filtered culture supernatants (Sup.) were analyzed by western blot.

**Extended Data Fig. 3.**
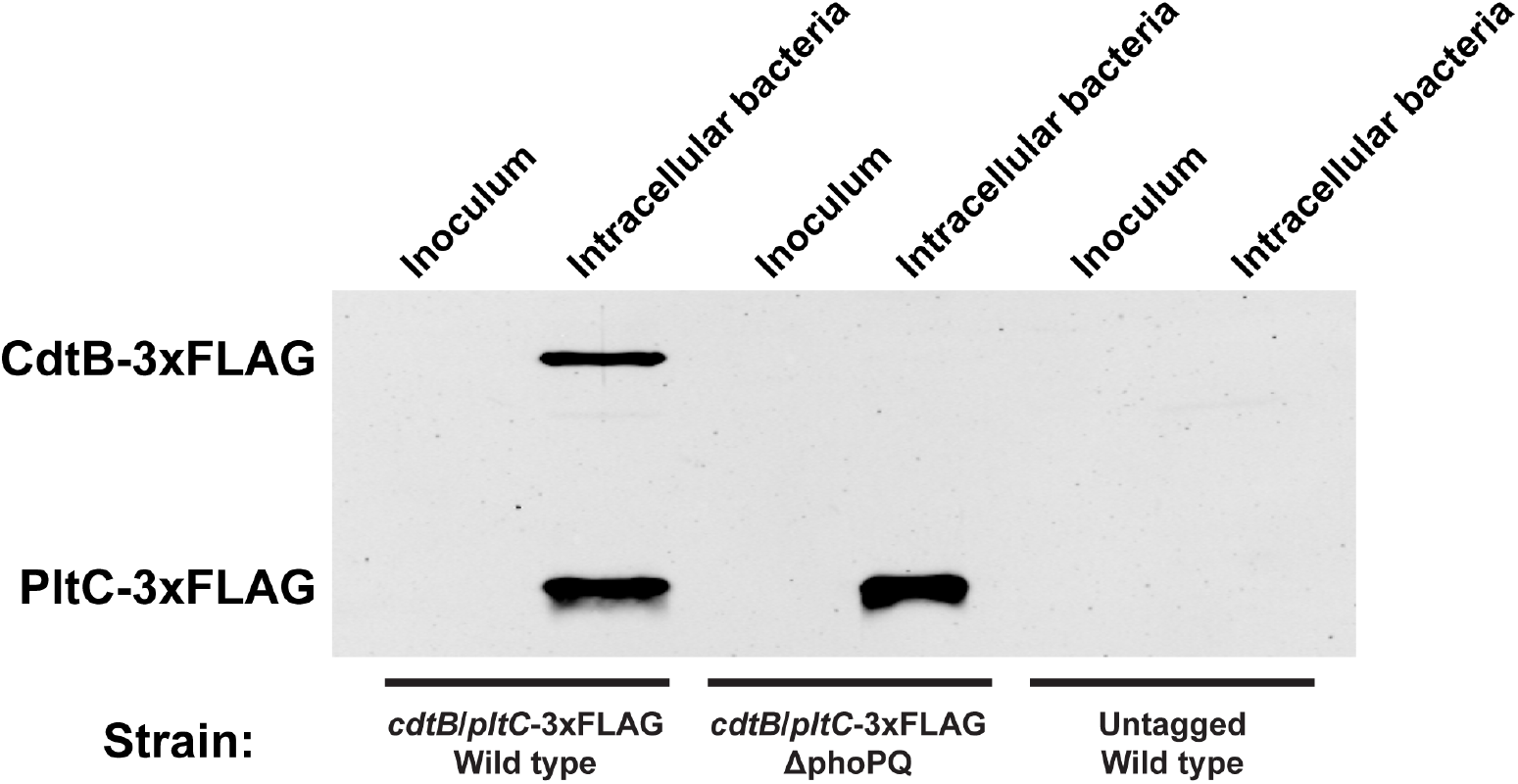
Expression of *pltC* by intracellular *S*. Typhi, unlike the other typhoid toxin genes, does not require the PhoP/PhoQ two-component system. Henle-407 cells were infected using wild type and Δ*phoP/phoQ* mutant *S*. Typhi strains that encode 3xFLAG epitope-tagged versions of both *cdtB* and *pltC* at their native genomic loci, as well as an untagged control strain. Whole cell lysates from the inoculum used for the infections as well as the bacteria isolated from infected cells 24 hours post-infection were analyzed by western blot. Sample loading was normalized to CFU recovery and lysate from 2.5×10^7^ bacteria was loaded in each lane.

**Extended Data Fig. 4.**
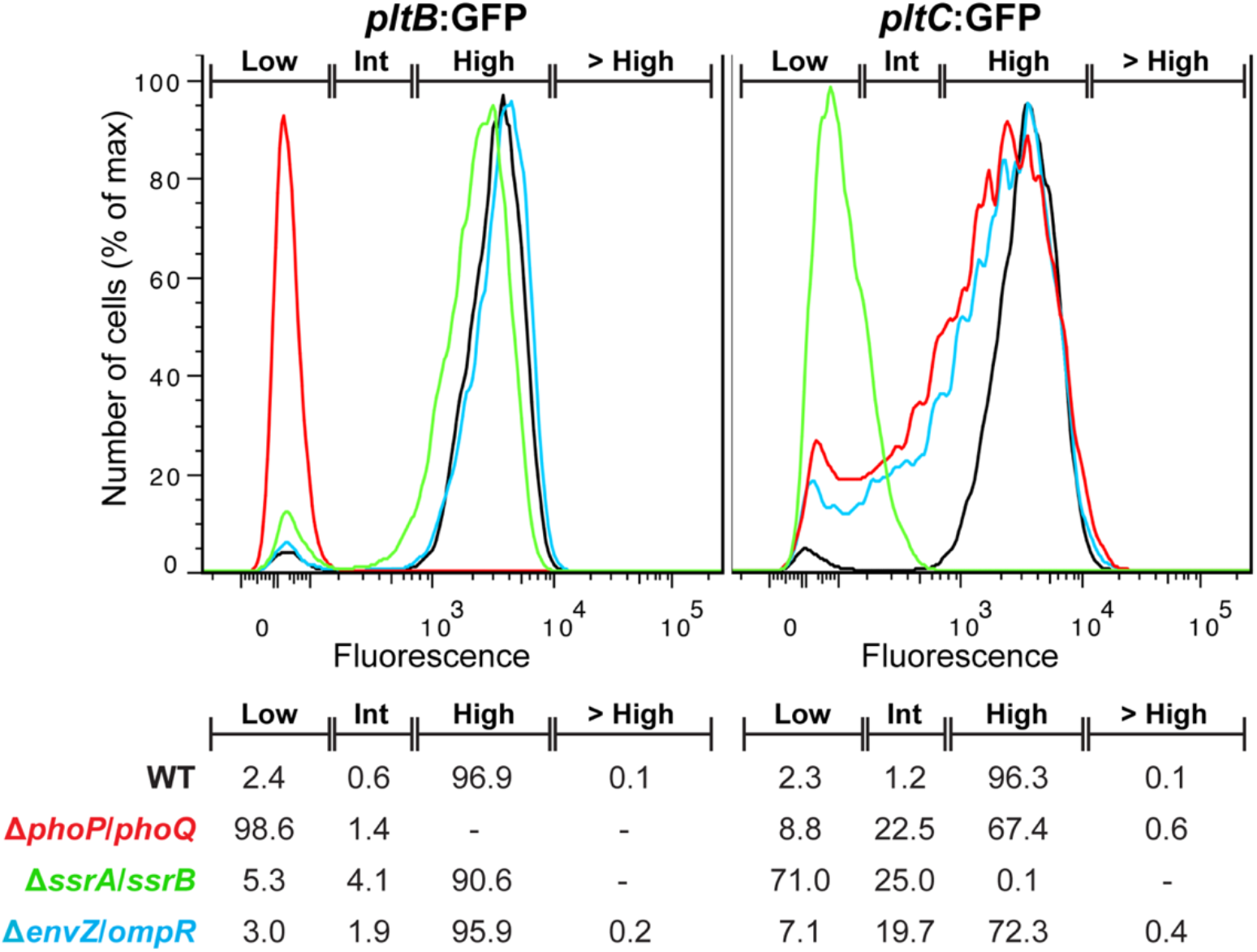
Single bacterium-level expression of the typhoid toxin B subunits by *S*. Typhi within infected cells in strains lacking key two-component regulators of intracellular gene expression. The SsrA/SsrB, PhoP/PhoQ and EnvZ/OmpR two-component regulatory systems are all central regulators of intracellular gene expression and all were identified as leading to reduced *pltC* expression in the FAST-INseq screen (Extended Data Table 1). To probe the role of these regulators in the expression of *pltB* and *pltC*, a flow cytometry analysis of *pltB:gfp* and *pltC:gfp* expression was conducted of the indicated *S*. Typhi strains 24 hours post-infection. Histograms show the GFP fluorescence intensities of individual bacteria. For both strains, the wild-type samples show high fluorescence (most bacteria) and low fluorescence (rare bacteria) populations; gates were established to show the percentage of bacteria in these populations for each strain, as well as those with an intermediate level of fluorescence (“int”) and those with a level greater than 99.9% of the wild-type population (> high). The proportions of bacteria with fluorescence intensities within these gates is shown (bottom). The heterogeneous expression of *pltC* in the Δ*phoP/phoQ* and Δ*envZ/ompR* strains is likely due to their influence over *ssrA/ssrB* expression ^24 36^.

**Extended Data Fig. 5.**
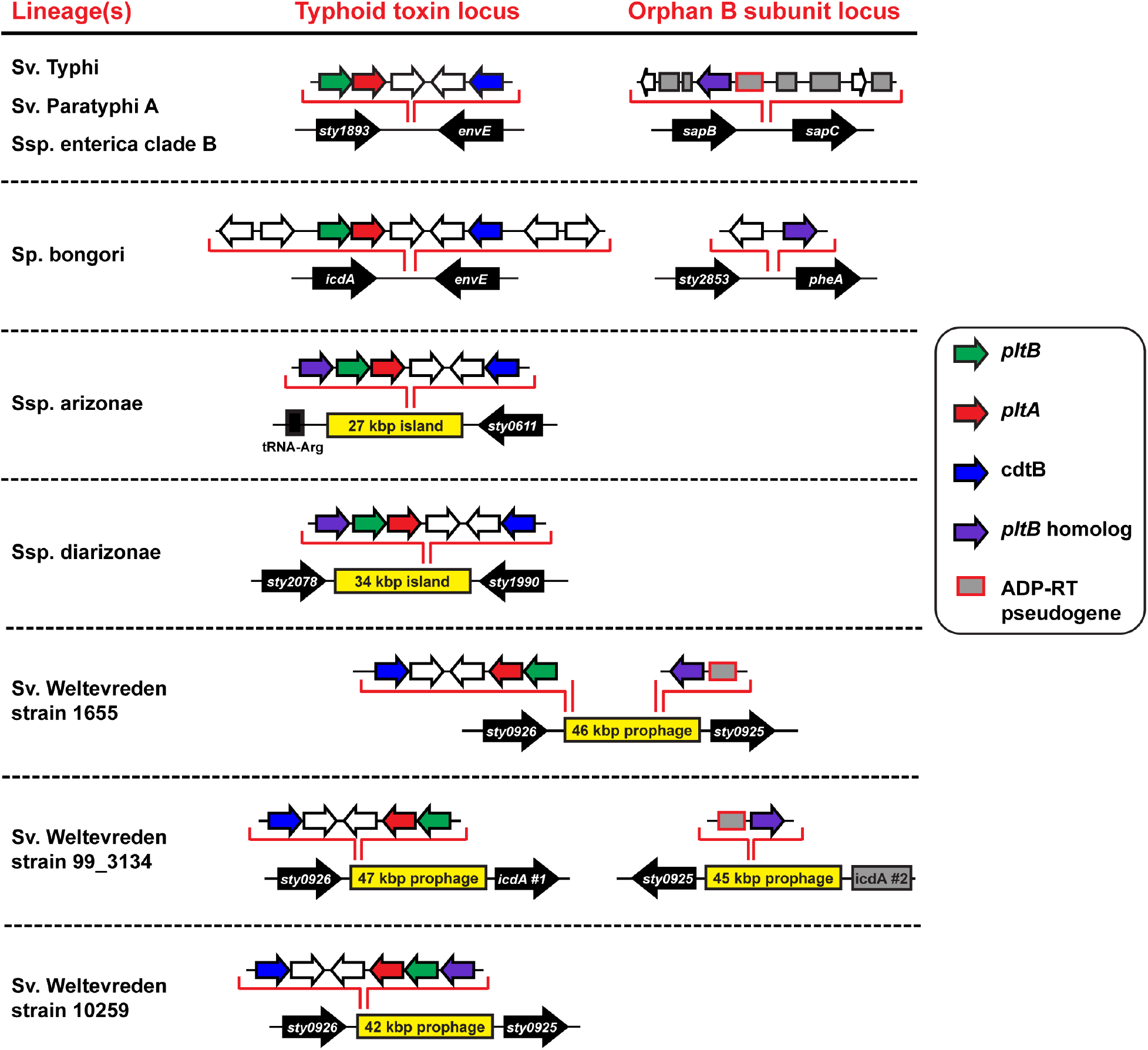
Congruent phylogenetic distributions of the typhoid toxin islet and a second pertussis toxin-like B subunit homolog in the *Salmonella* genus. Depiction of different genomic arrangements of the typhoid toxin islet and of loci encoding a second putative B subunit that were identified by searching the NCBI genome database. All genomic locations are shown relative to the *S*. Typhi reference genome (black arrows). Large genomic islands absent from the Typhi genome are shown in yellow, pseudogenes are shown as grey boxes and ADP-ribosyltransferase (ADP-RT) pseudogenes have a red outline. The depicted genomic arrangements appear, based on the available sequence data, to be conserved across the cited species (Sp.), subspecies (Ssp.), clade, serovar (Sv.) or strains, as indicated.

**Extended Data Fig. 6.**
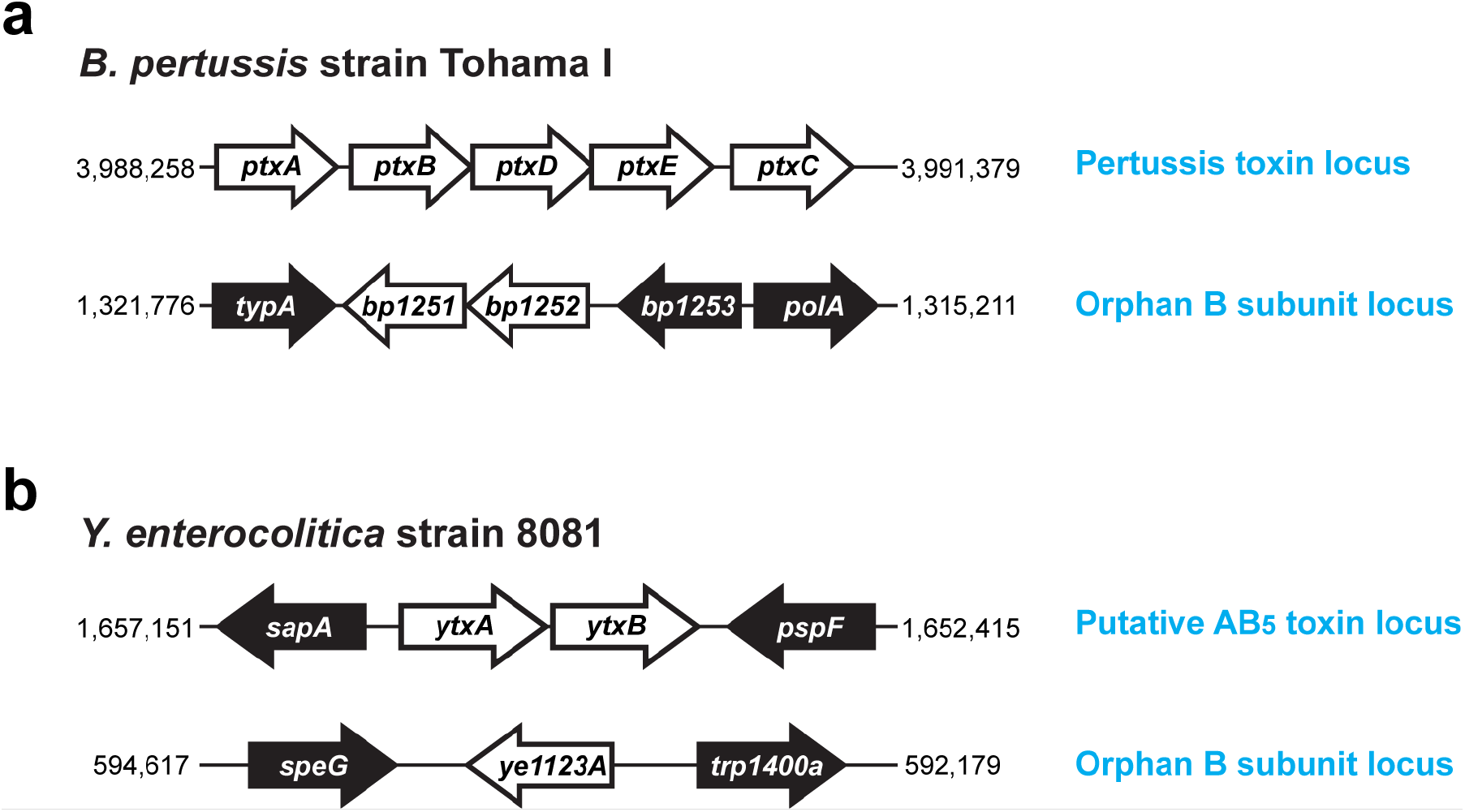
Examples of AB_5_-type toxins and orphan homologous delivery subunits encoded within the same genome outside the *Salmonella* genus. Using BLAST homology searches using AB_5_-type toxin subunits, genomes encoding an AB_5_-type toxin as well as a homologous orphan delivery subunit were identified and are depicted above. (**a**) In addition to the five-gene pertussis toxin locus ^37^, we found that *B. pertussis* encodes two putative AB_5_ toxin delivery subunits at a distant genomic locus that does not also encode a putative toxin active subunit, *bp1251* and *bp1252*. Bp1251 is >30% identical to both PtxB and PtxC over >200 amino acids. Bp1252 was identified as a statistically significant hit (e value < 0.02) to PtxB, PtxC and PtxE using the HHPred protein homology detection tool (https://toolkit.tuebingen.mpg.de/#/tools/hhpred). (**b**) The putative AB_5_-type toxin Yersinia toxin ^38^ is encoded by *ytxA* and *ytxB* in *Y. enterocolitica*. At a distant genomic location, *Y. enterocolitica* also encodes an orphan pertussis toxin-like delivery subunit, *ye1123A*. YtxA and Ye1123A exhibit significant sequence similarity to the pertussis-like toxin ArtAB; YtxA is > 60% identical to ArtA and Ye1123A is ~30% identical to ArtB.

**Extended Data Table 1:** Genes identified by FAST-INSeq as important for *pltC* expression in infected human cells.

| Gene | High fluor:<br>insertions | High fluor:<br>reads | Low fluor:<br>insertions | Low fluor:<br>reads | Ratio<sup>a</sup> |
| --- | --- | --- | --- | --- | --- |
| Gene regulation |  |  |  |  |  |
| ssrB | 22 | 48 | 27 | 936 | 20 |
| ssrA | 36 | 52 | 61 | 839 | 16 |
| phoP | 13 | 33 | 15 | 201 | 6 |
| phoQ | 26 | 62 | 27 | 335 | 5.5 |
| ompR | 14 | 73 | 13 | 244 | 3.5 |
| Pyrimidine biosynthesis |  |  |  |  |  |
| carA | 10 | 98 | 11 | 897 | 9 |
| carB | 35 | 194 | 33 | 1669 | 8.5 |
| pyrD | 21 | 139 | 20 | 979 | 7 |
| pyrE | 19 | 477 | 16 | 2705 | 5.5 |
| pyrB | 19 | 122 | 16 | 536 | 4.5 |
| pyrF | 10 | 40 | 7 | 188 | 4.5 |
| Other |  |  |  |  |  |
| proB | 20 | 307 | 20 | 2449 | 8 |
| proC | 12 | 216 | 11 | 1343 | 6 |
| rplS | 3 | 45 | 3 | 238 | 5.5 |
| proA | 27 | 798 | 24 | 3704 | 4.5 |
| ugpQ | 15 | 677 | 13 | 2652 | 4 |
| t3900 (bcsZ) | 25 | 260 | 19 | 864 | 3.5 |
| lysA | 33 | 1004 | 30 | 3497 | 3.5 |
| t4432 (ulaR) | 16 | 86 | 8 | 308 | 3.5 |
| vexA | 33 | 1099 | 25 | 3216 | 3 |
| t3052 | 13 | 549 | 10 | 1292 | 2.5 |
| tviB | 45 | 1990 | 35 | 5021 | 2.5 |

<sup>a</sup>: Ratio of normalized INSeq sequencing reads from transposon insertions within the indicated gene in the low fluorescence pool compared to the high fluorescence pool.

**Extended Data Table 2:**
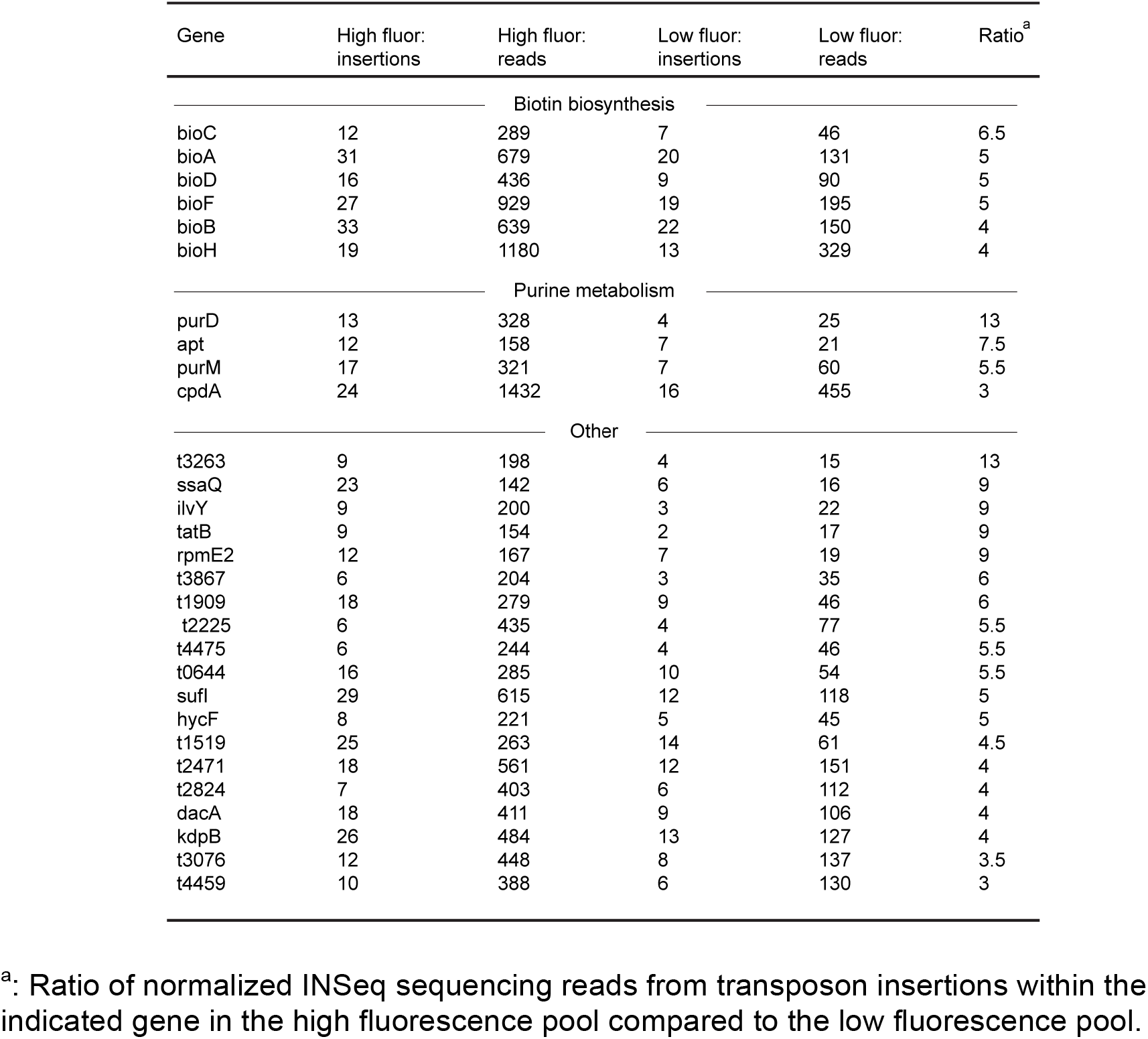
Mutants identified by FAST-INSeq that lead to an increased fraction of *S*. Typhi expressing high levels of *pltC* in infected human cells.

| Gene | High fluor:<br>insertions | High fluor:<br>reads | Low fluor:<br>insertions | Low fluor:<br>reads | Ratio<sup>a</sup> |
| --- | --- | --- | --- | --- | --- |
| Biotin biosynthesis |  |  |  |  |  |
| bioC | 12 | 289 | 7 | 46 | 6.5 |
| bioA | 31 | 679 | 20 | 131 | 5 |
| bioD | 16 | 436 | 9 | 90 | 5 |
| bioF | 27 | 929 | 19 | 195 | 5 |
| bioB | 33 | 639 | 22 | 150 | 4 |
| bioH | 19 | 1180 | 13 | 329 | 4 |
| Purine metabolism |  |  |  |  |  |
| purD | 13 | 328 | 4 | 25 | 13 |
| apt | 12 | 158 | 7 | 21 | 7.5 |
| purM | 17 | 321 | 7 | 60 | 5.5 |
| cpdA | 24 | 1432 | 16 | 455 | 3 |
| Other |  |  |  |  |  |
| t3263 | 9 | 198 | 4 | 15 | 13 |
| ssaQ | 23 | 142 | 6 | 16 | 9 |
| ilvY | 9 | 200 | 3 | 22 | 9 |
| tatB | 9 | 154 | 2 | 17 | 9 |
| rpmE2 | 12 | 167 | 7 | 19 | 9 |
| t3867 | 6 | 204 | 3 | 35 | 6 |
| t1909 | 18 | 279 | 9 | 46 | 6 |
| t2225 | 6 | 435 | 4 | 77 | 5.5 |
| t4475 | 6 | 244 | 4 | 46 | 5.5 |
| t0644 | 16 | 285 | 10 | 54 | 5.5 |
| sufI | 29 | 615 | 12 | 118 | 5 |
| hycF | 8 | 221 | 5 | 45 | 5 |
| t1519 | 25 | 263 | 14 | 61 | 4.5 |
| t2471 | 18 | 561 | 12 | 151 | 4 |
| t2824 | 7 | 403 | 6 | 112 | 4 |
| dacA | 18 | 411 | 9 | 106 | 4 |
| kdpB | 26 | 484 | 13 | 127 | 4 |
| t3076 | 12 | 448 | 8 | 137 | 3.5 |
| t4459 | 10 | 388 | 6 | 130 | 3 |

<sup>a</sup>: Ratio of normalized INSeq sequencing reads from transposon insertions within the indicated gene in the high fluorescence pool compared to the low fluorescence pool.

## Reference

1 Dougan, G. & Baker, S. Salmonella enterica serovar Typhi and the pathogenesis of typhoid fever. Annu Rev Microbiol 68, 317–336, doi:10.1146/annurev-micro-091313-103739 (2014).

2 Parry, C. M., Hien, T. T., Dougan, G., White, N. J. & Farrar, J. J. Typhoid fever. N Engl J Med 347, 1770–1782, doi:10.1056/NEJMra020201 (2002).

3 Spano, S., Ugalde, J. E. & Galan, J. E. Delivery of a Salmonella Typhi exotoxin from a host intracellular compartment. Cell Host Microbe 3, 30–38, doi:10.1016/j.chom.2007.11.001 (2008).

4 Song, J., Gao, X. & Galan, J. E. Structure and function of the Salmonella Typhi chimaeric A(2)B(5) typhoid toxin. Nature 499, 350–354, doi:10.1038/nature12377 (2013).

5 Galan, J. E. Typhoid toxin provides a window into typhoid fever and the biology of Salmonella Typhi. Proc Natl Acad Sci U S A 113, 6338–6344, doi:10.1073/pnas.1606335113 (2016).

6 Gao, X. et al. Evolution of host adaptation in the Salmonella typhoid toxin. Nat Microbiol. 2, 1592–1599 (2017).

7 Haghjoo, E. & Galan, J. E. Salmonella typhi encodes a functional cytolethal distending toxin that is delivered into host cells by a bacterial-internalization pathway. Proc Natl Acad Sci U S A 101, 4614–4619, doi:10.1073/pnas.0400932101 (2004).

8 Fowler, C. & Galán, J. Decoding a Salmonella Typhi Regulatory Network that Controls Typhoid Toxin Expression within Human Cells. Cell Host Microbe 23, 65–76 (2018).

9 Merritt, E. & Hol, W. AB5 toxins. Curr Opin Struct Biol. 5, 165–171 (1995).

10 Stein, P. et al. The crystal structure of pertussis toxin. Structure 2, 45–57 (1994).

11 Locht, C., Coutte, L. & Mielcarek, N. The ins and outs of pertussis toxin. FEBS J. 278, 4668–4682 (2011).

12 Lara-Tejero, M. & Galan, J. E. A bacterial toxin that controls cell cycle progression as a deoxyribonuclease I-like protein. Science 290, 354–357 (2000).

13 Geiger, T., Pazos, M., Lara-Tejero, M., Vollmer, W. & Galán, J. Peptidoglycan editing by a specific LD-transpeptidase controls the muramidase-dependent secretion of typhoid toxin. Nat Microbiol. 3, 1243–1254 (2018).

14 Chang, S., Song, J. & Galán, J. Receptor-Mediated Sorting of Typhoid Toxin during Its Export from Salmonella Typhi-Infected Cells. Cell Host Microbe 20, 682–689 (2016).

15 Song, J., Gao, X. & Galan, J. E. Structure and function of the Salmonella Typhi chimaeric A(2)B(5) typhoid toxin. Nature 499, 350–354 (2013).

16 Gawronski, J. D., Wong, S. M., Giannoukos, G., Ward, D. V. & Akerley, B. J. Tracking insertion mutants within libraries by deep sequencing and a genome-wide screen for Haemophilus genes required in the lung. Proc Natl Acad Sci U S A 106, 16422–16427, doi:10.1073/pnas.0906627106 (2009).

17 Goodman, A. L. et al. Identifying genetic determinants needed to establish a human gut symbiont in its habitat. Cell Host Microbe 6, 279–289, doi:10.1016/j.chom.2009.08.003 (2009).

18 van Opijnen, T., Bodi, K. L. & Camilli, A. Tn-seq: high-throughput parallel sequencing for fitness and genetic interaction studies in microorganisms. Nat Methods 6, 767–772, doi:10.1038/nmeth.1377 (2009).

19 Ochman, H., Soncini, F. C., Solomon, F. & Groisman, E. A. Identification Of a Pathogenicity Island Required For Salmonella Survival In Host Cells. Proc. Natl. Acad. Sc. USA 93, 7800–7804 (1996).

20 Shea, J. E., Hensel, M., Gleeson, C. & Holden, D. W. Identification of a Virulence Locus Encoding a Second Type III Secretion System in *Salmonella Typhimurium*. Proc. Natl. Acad. Sc. USA 93, 2593–2597 (1996).

21 Hensel, M. et al. Genes encoding putative effector proteins of the type III secretion system of Salmonella pathogenicity island 2 are required for bacterial virulence and proliferation in macrophages. Mol Microbiol. 30, 163–174 (1998).

22 Cirillo, D., Valdivia, R., Monack, D. & Falkow, S. Macrophage-dependent induction of the Salmonella pathogenicity island 2 type III secretion system and its role in intracellular survival. Mol. Microbiol. 30, 175–188 (1998).

23 Garmendia, J., Beuzón, C., Ruiz-Albert, J. & Holden, D. The roles of SsrA-SsrB and OmpR-EnvZ in the regulation of genes encoding the Salmonella typhimurium SPI-2 type III secretion system. Microbiology 149, 2385–2396 (2003).

24 Fass, E. & Groisman, E. Control of Salmonella pathogenicity island-2 gene expression. Curr Opin Microbiol. 12, 199–204 (2009).

25 Miller, R. et al. The Typhoid Toxin Produced by the Nontyphoidal Salmonella enterica Serotype Javiana Is Required for Induction of a DNA Damage Response In Vitro and Systemic Spread In Vivo. MBio 9, pii: e00467–00418. (2018).

26 Miller, R. & Wiedmann, M. The Cytolethal Distending Toxin Produced by Nontyphoidal Salmonella Serotypes Javiana, Montevideo, Oranienburg, and Mississippi Induces DNA Damage in a Manner Similar to That of Serotype Typhi. MBio 7, pii: e02109–02116. (2016).

27 Rodriguez-Rivera, L., Bowen, B., den Bakker, H., Duhamel, G. & Wiedmann, M. Characterization of the cytolethal distending toxin (typhoid toxin) in non-typhoidal Salmonella serovars. Gut Pathog. 7, 19 (2015).

28 Luu, L. et al. Characterisation of the Bordetella pertussis secretome under different media. J Proteomics 158, 43–51 (2017).

29 Galán, J. E. & Curtiss III, R. Distribution of the invA, -B, -C, and -D genes of Salmonella typhimurium among other Salmonella serovars: invA mutants of Salmonella typhi are deficient for entry into mammalian cells. Infect. Immun. 59, 2901–2908 (1991).

30 Demarre, G. et al. A new family of mobilizable suicide plasmids based on broad host range R388 plasmid (IncW) and RP4 plasmid (IncPα) conjugative machineries and their cognate Escherichia coli host strains. Res. Microbiol. 156, 245–255 (2005).

31 Snavely, M., Miller, C. & Maguire, M. The mgtB Mg2+ transport locus of Salmonella typhimurium encodes a P-type ATPase. J Biol Chem. 266, 815–823 (1991).

32 Miller, J. H. Experiments in molecular genetics. (Cold Spring Harbor Laboratory, 1972).

33 Spanò, S., Gao, X., Hannemann, S., Lara-Tejero, M. & Galán, J. A Bacterial Pathogen Targets a Host Rab-Family GTPase Defense Pathway with a GAP. Cell Host Microbe 19, 216–226 (2016).

34 van Opijnen, T., Bodi, K. & Camilli, A. Tn-seq: high-throughput parallel sequencing for fitness and genetic interaction studies in microorganisms. Nat Methods. 6, 767–772 (2009).

35 Goodman, A. et al. Identifying genetic determinants needed to establish a human gut symbiont in its habitat. Cell Host Microbe. 6, 279–289 (2009).

36 Lee, A., Detweiler, C. & Falkow, S. OmpR regulates the two-component system SsrA-ssrB in Salmonella pathogenicity island 2. J. Bacteriol. 182, 771–781 (2000).

37 Gross, R., Aricò, B. & Rappuoli, R. Genetics of pertussis toxin. Mol Microbiol. 3, 119–124 (1989).

38 Axler-Diperte, G., Miller, V. & Darwin, A. YtxR, a conserved LysR-like regulator that induces expression of genes encoding a putative ADP-ribosyltransferase toxin homologue in Yersinia enterocolitica. J Bacteriol. 188, 8033–8043 (2006).

